# LIKE EARLY STARVATION is involved in the regulation of starch initiation in potato (*Solanum tuberosum* cv. Désirée) tubers

**DOI:** 10.1101/2025.06.23.661224

**Authors:** Camille Locquet, Mara Berti, Fabrice Bray, Hyazann Hulin, Gregory Stoclet, David Seung, Denis Lourdin, Christophe D’Hulst, Fabrice Wattebled, Nicolas Szydlowski

## Abstract

Starch initiation has been intensively investigated. In tubers of *Solanum tuberosum*, it is controlled by a heteromultimeric isoamylase complex composed of ISA1 and ISA2. Moreover, two non-catalytic starch binding proteins (i.e., ESV1 and LESV) regulate leaf starch metabolism in Arabidopsis. Both proteins interact with starch glucans through a C-terminal antiparallel β-sheet domain and differ by the presence of a 115 amino acid N-terminal overhang in LESV. Here we report the CRIPSR/Cas9 inactivation of *LESV* or *ESV1* in potato and the characterization of the respective tuber starches. Starch from *esv1* mutants was unaffected whereas granule diameter was severely reduced in tubers of *lesv* plants. This was negatively correlated to the number of granules, coherent with an altered starch initiation phenotype. Strikingly, scanning electron microscopy of purified starches revealed that *lesv* plants phenocopy *isa1* and *isa2* antisense lines described 20 years ago. A physical interaction between LESV and ISA1 was previously demonstrated in rice. Here, we confirmed this interaction in potato using yeast two-hybrid assays, providing molecular evidence for their functional association in potato. This interaction supports the hypothesis that LESV acts in concert with the ISA1/ISA2 complex, likely regulating the early steps of glucan organization required for proper starch granule formation.

## Introduction

Starch is composed of two glucose polymers, amylose and amylopectin, the latter representing about 80% of the starch weight in potato (Fredriksson *et al*., 1998). Amylose, essentially linear, consists of glucose residues linked by α-1,4 O-glycosidic bonds whereas amylopectin is a branched glucan characterized by glucose residues linked by α-1,4 O-glycosidic bonds with branching points formed by α-1,6 O-glycosidic bonds connecting the linear chains together. In potato tuber, amylose and amylopectin arrange to form a single large semicrystalline granule within the amyloplast (Kram *et al*., 1993). Despite the apparent simplicity of its structure, merely consisting of glucose residues and two types of bonds, starch metabolism is in fact a complex process involving a wide range of proteins. Amylose is mainly synthesized by the Granule Bound Starch Synthase (GBSS), which is the most abundant starch-bound protein (Vos-Scheperkeuter *et al*., 1986; Ball *et al*., 1998). In contrast, the synthesis of amylopectin involves a diverse array of enzymes that, beyond the ADP-glucose pyrophosphorylase (AGPase) which is shared with amylose synthesis for generating the ADP-Glucose precursor, include soluble starch synthases (SSs), starch branching enzymes (BEs), starch debranching enzymes (DBEs) and starch dikinases (GWD and PWD) (Reyniers *et al*., 2020).

Starch initiation in plants has been deeply investigated in recent years. Due to the availability of efficient molecular toolboxes to study this process, an important focus was put on transitory starch from the model plant *Arabidopsis thaliana* (Seung & Smith, 2019). In the last two decades, several proteins were demonstrated to participate in the granule initiation process in chloroplasts (Seung & Smith, 2019; Mérida & Fettke, 2021; Vandromme *et al*., 2023; Sharma *et al*., 2024). Protein targeting to starch 2 (PTST2) binds both to soluble maltooligosaccharide (MOS) primers and starch synthase 4 (SS4) (Seung *et al*., 2017). The latter elongates MOSs further, prior to the action of the other synthesizing enzymes (i.e., SSs, BEs and DBEs) (Roldán *et al*., 2007; Seung *et al*., 2016; Lu *et al*., 2018). This protein/glucan initiation complex also includes MRC/Pii1 (myosin-resembling chloroplast protein/protein involved in starch initiation 1). PTST2 is recruited by the thylakoid-bound protein MFP1 (mar-binding filament-like protein 1) to specific areas of the stroma, in the vicinity of thylakoids, where it develops starch granules (Seung *et al*., 2018; Vandromme *et al*., 2019; Sharma *et al*., 2024). Starch initiation in amyloplasts may involve some of the aforementioned proteins, though the function of SS4 remains unclear, and that concentration/location of the initiation components most likely differs from chloroplasts due to the absence of thylakoids (Satoh *et al*., 2008; Toyosawa *et al*., 2016; Jung *et al*., 2018; Seung & Smith, 2019).

For potato tubers, altered starch initiation has only been reported for lines with antisense inhibition of the debranching enzymes, isoamylase 1 (ISA1) or 2 (ISA2) (Bustos *et al*., 2004). Both proteins form a heteromultimeric complex in all species, although ISA1 can also act alone as homomultimer in some species, such as maize or rice (Hussain *et al*., 2003; Utsumi & Nakamura, 2006; Kubo *et al*., 2010; Facon *et al*., 2013). ISA1/ISA2 complex is responsible for hydrolyzing some of the α-1,6 linkages to organize a distribution of branch points compatible with amylopectin phase transition (Ball *et al*., 1996). Inactivation of the complex in Arabidopsis leaves and cereal endosperm leads to the accumulation of branched soluble glucans called “phytoglycogen” (Delatte *et al*., 2005; Wattebled *et al*., 2005, 2008; Streb *et al*., 2009). In potato, reduced ISA1/ISA2 activity leads to more limited phytoglycogen accumulation in some antisense lines, but a large increase of granule numbers per amyloplast, and strong decrease in starch granule diameter, suggesting an impaired regulation of starch initiation (Bustos *et al*., 2004). The proposed mechanism relies on the capacity of ISA1/ISA2 to regulate the number of starch granules per amyloplast by limiting the initiation of new glucans precursors.

Recent studies in *Arabidopsis thaliana* revealed two non-catalytic proteins, ESV1 (early starvation 1) and LESV (like early starvation 1). Despite the lack of known catalytic domains, ESV1 plays a crucial role in regulating transitory starch degradation in *Arabidopsis* leaves (Feike *et al*., 2016). Starch turnover is impaired in *esv1* mutant rosettes with faster starch degradation at night, whereas no modifications of the starch content over the day/night cycle was observed in *lesv* mutants. On the other hand, LESV-overexpressing plants exhibited a 3-fold decrease of the starch amount during the night. Granules from *esv1* and *lesv* mutants displayed irregular shapes compared to wild-type starch. On the other hand, starch granule size was decreased or increased in LESV- or ESV1-overexpressors, respectively. In LESV-overexpressing plants, the number of starch granules per chloroplast was also increased. Furthermore, recombinant Arabidopsis ESV1 and LESV decreased the activity of GWD and starch phosphorylase (PHS1), or enhanced the activity of SS1, SS3, β-amylase and isoamylase on starch granules *in vitro* (Singh *et al*., 2022). LESV and ESV1 share a highly similar C-terminal region organized in an antiparallel β-sheet domain which harbors alignments of Trp residues. To date, all the reports on these proteins showed physical interactions with starch granules, which were proposed to rely on these aligned residues (Feike *et al*., 2016; Singh *et al*., 2022; Liu *et al*., 2023; Yan *et al*., 2024; Osman *et al*., 2024). Both recombinant proteins from Arabidopsis were shown to interact with amylopectin in electrophoretic mobility shift assays and SR-CD analysis (Osman *et al*., 2024). Interestingly, the LESV amino-acid sequence contains an N-terminal extension that is absent from ESV1, and the conformation of this extension is influenced by amylopectin binding *in vitro* (Osman *et al*., 2024). By combining synthetic biology in yeast, structural biology and functional genomics in Arabidopsis, it was recently proposed that LESV and ESV1 promote and stabilize amylopectin phase transition, respectively (Liu *et al*., 2023). The authors established a model in which LESV promotes the amylopectin phase transition by helping align double helices, and ESV1 stabilizes the formed semi-crystalline lamellae.

More recently, the rice (*Oryza sativa*) *flo9* mutant, harboring a phenotype similar to that of *isa1* mutants (*i.e.*, defect in amylopectin synthesis and a floury endosperm), was shown to be deficient in the ortholog of Arabidopsis *LESV* (Yan *et al*., 2024). Yeast two-hybrid, *in vitro* pulldown and *in vivo* firefly luciferase complementation assays revealed direct interaction between rice LESV and ISA1 polypeptides. This was attributed to the N-terminus region of LESV by domain truncation analysis. In this study, interaction between LESV and PTST2 was also reported and both proteins were proposed to act synergistically as cargos for controlling starch biosynthesis and endosperm development by targeting ISA1 to starch. In a previous work, we identified LESV as one of the major starch-binding proteins after GBSS in potato tubers by nanoLC-nanoESI-MS/MS (Helle *et al*., 2018). ESV1 was also observed in this analysis, but with a much lower abundance. Interestingly, both proteins were unaffected by extended detergent or proteolytic shaving of the starch granule surface. Postulating that their presence within the granule illustrates a role in tuber starch metabolism, we used CRISPR/Cas9 genome editing to investigate their function in potato.

Here, we report on the production of the corresponding knock-out mutant lines and their phenotypic analysis. While *esv1* mutants did not show any change in granule size and shape, *lesv* lines displayed a dramatic decrease of granule diameter and the accumulation of small granule aggregates, indicating deregulated starch initiation. Detailed characterization of starch ultra structure did not reveal any change in both mutants apart from an increase in phosphate contents and the propensity to amylase digestibility in *lesv* mutants, likely as a consequence of reduced average granule diameter. Strikingly, *lesv* phenotype is a phenocopy of *isa1* and *isa2* antisense potato lines described by Bustos and collaborators in 2004 (Bustos *et al*., 2004). In the present study, we confirmed by yeast two-hybrid assay that, similar to the rice proteins, potato LESV and ISA1 physically interact. Finally, the absence of phenotype in *esv1* mutants highlights the critical role of the LESV-specific interaction in the regulation of starch initiation. Our data suggest that *LESV* regulates starch initiation in potato tubers by targeting ISA1 to new granule precursors, which, in turn, controls the number of starch granules per amyloplast by restricting the number of initiation events.

## Results

### 1. CRISPR/Cas9 inactivation of *LESV* and *ESV1* has no impact on plant development, starch and phytoglycogen contents

Targeted mutagenesis in potato faces challenges due to its tetraploid complex genomic structure. To overcome this, we used the CRISPR/Cas9 genome editing approach, which offers the ability to target all four alleles simultaneously with high efficiency and specificity. Guide RNAs (gRNAs) specific to the four copies of *ESV1* and *LESV* in Désirée were designed using Geneious software, along with CRISPR-P and CRISPOR online tools as well as the polymorphism map described in (Sevestre *et al*., 2020). All four copies of *ESV1* and *LESV* shared identical sequences at the selected gRNAs that targeted the third exon in both genes (Fig. 1a and 1b). Subsequently, gRNAs were cloned by Golden Gate reaction into the pDIRECT_22A plasmid, which carries a kanamycin resistance gene (Čermák *et al*., 2017) (Supplemental Fig.S1). This construct was then delivered into potato plants via *Agrobacterium tumefaciens* nuclear genome transformation. To confirm that transformation was successful, the presence of the *CAS9* gene in the genome was screened by PCR amplification using specific primers (Table S1 and Fig. 1c). Genotyping of the regenerated Kan^R^ plants was achieved by sequencing the targeted regions of *LESV* and *ESV1* after amplification by PCR, with the primers listed in Table S1. Sequence chromatograms were analyzed with the ICE analysis tool (https://ice.synthego.com/<u>#/</u>) (Supplemental Fig. S2 and S3). ICE Synthego analysis separates overlapping signals corresponding to different alleles. The number of edited alleles was deducted from the inferred sequences and their relative proportions, considering that one allele corresponds to approximately 25% of the signal. Among 35 and 31 transformants, confirmed by PCR amplification of the *CAS9* gene, 28 and 21 had at least one mutated allele for *LESV* and ESV1 respectively. Nine independent full knock-out lines were selected for *LESV* and five for *ESV1* (Fig. 1d). This corresponded to 32.2% and 23.8% of the transformants, respectively. In the selected lines, all four alleles contained frameshift mutations (Fig. 1d). Granule morphology was investigated in all generated lines and subsequent analyses were performed on three or two selected *lesv* and *esv1* lines (Fig. 1d). The absence of the corresponding proteins in mutant starches was confirmed by label-free quantitative proteomics following starch gelatinization and protein isolation (supplemental Fig. S4). Mutants were cultivated on soil for three months with a photoperiod of 16 h light (80 µmol.m^−2^.s^−1^) at 22°C /8 h of darkness at 20°C and various parameters were assessed (Fig. 2). No significant modifications were observed in the overall appearance of plants, stem height, and tuber morphology of both mutants (Fig. 2a to 2f). Furthermore, both tuber yield and number per plant remained unaffected by the mutation (Fig. 2g and 2h). In addition, starch, glucose and water-soluble polysaccharides (WSPs) contents were measured (Fig. 2i and 2j). No consistent changes could be observed across all *lesv* lines (Fig. 2i). In *esv1*, both lines had more WSPs, but the overall amount was low (ie., <10% of total starch content) (Fig. 2j).

**Figure 1:**
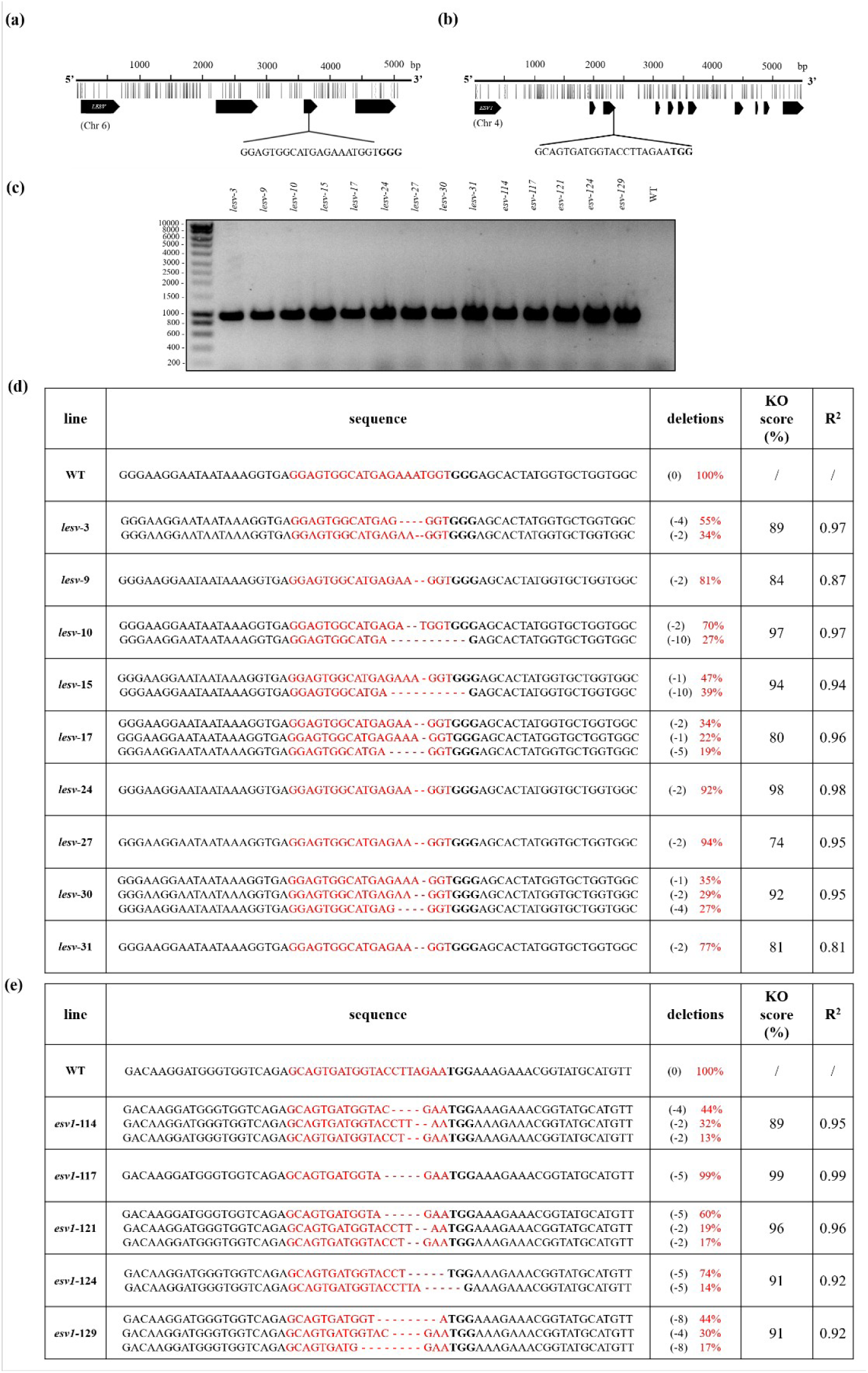
Targeted mutagenesis of the *LESV* and ESV1 genes using CRISPR/Cas9 system. (a, b) Target site of the guide RNAs (gRNAs) within the *LESV* and *ESV1* genes, respectively. Black arrows represent exons, and the Protospacer Adjacent Motif (PAM) is highlighted in bold. Vertical lines above exons are single nucleotide polymorphisms (transversions), broken lines are insertions. Chr: chromosome. (c) PCR amplification of a 959 bp region of the *CAS9* gene in mutants. The first lane corresponds to the molecular weight standard, in bp. The last lane is a negative control, from a wild type (WT) plant. (d, e) Genotyping results of the nine *lesv* mutants and five *esv1* mutants, identified as shift mutants using the ICE Synthego algorithm. In the sequences, red letters highlight the gRNA and the PAM motif is marked in bold. The “deletion” column specifies the number of deleted bases (in parentheses) in the inferred sequences, with their relative proportions in the edited population noted in red. The KO (knock-out) score is the proportion of deletions that led to a frameshift, the R^2^ value indicates how well the proposed deletions fit the Sanger sequence data of the edited sample. Differences between the percentages in the “deletions” column and the KO score are attributed to weak signals (∼5% of the edited population, not represented here), potentially coming from PCR amplification or sequencing errors, or misinterpretation by the Synthego algorithm.

**Figure 2:**
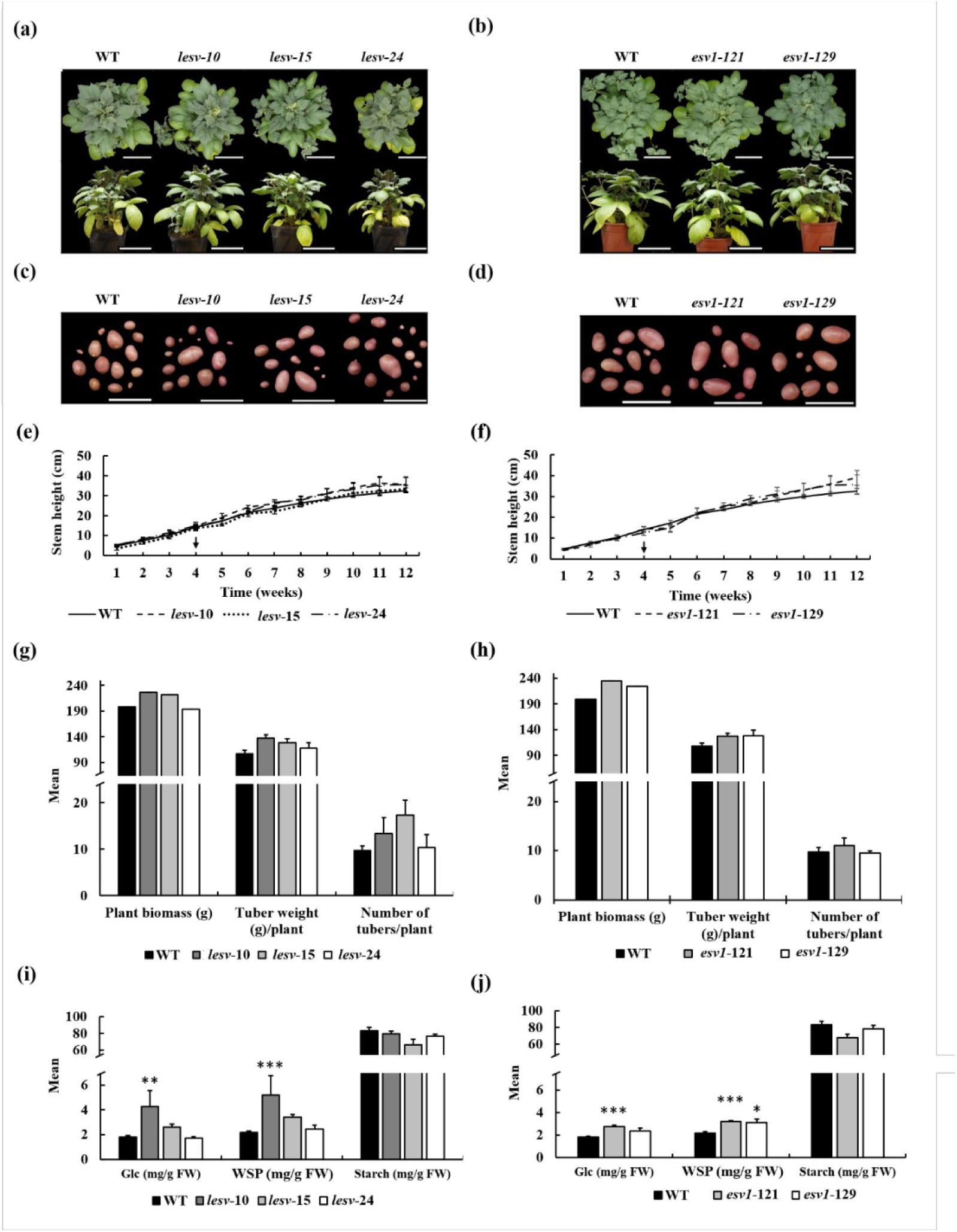
Potato growth and development. (a, b) Pictures of two-month-old wild type (WT) and mutant plants grown on soil in a growth cabinet with 16 h light/8 h dark photoperiod (80 µmol.m^−2^.s^−1^). (c, d) Tubers harvested from three-month-old WT or *lesv* and *esv1* mutant plants. (e, f) Stem size of WT and mutant plants over 12 weeks of culture. The black arrow indicates the moment when additional soil was added to the base of the stem. For WT, values are the mean (+/− SE) of three independent biological replicates from three independent cultures (n=9). For mutant lines, values are the mean (+/− SE) of three independent cultures (n=3). (g, h) Plant biomass, tuber weight per plant, and tuber number per plant of three-month-old plants. Except for plant biomass that was measured in a unique culture, values are mean (+/− SE) of three independent cultures. (i, j) Glucose (Glc), water-soluble polysaccharides (WSPs) and starch contents in tubers expressed in mg.g^−1^ of fresh weight (FW). For WT, values are mean (+/− SE) of three biological replicates from three independent cultures and three experimental replicates for each culture (n=27). For each mutant line, values are mean (+/− SE) of three independent cultures and three experimental replicates for each culture (n=9). Statistical analysis was made using Mann-Whitney tests, comparing the mutant line to the wild type (* = *p* ≤ 0,05; ** = *p* ≤ 0,01; *** = *p* ≤ 0,005). Scale bar in (a) and (b) = 15 cm, and in (c) and (d) = 10 cm.

### 2. Starch granule morphology is drastically affected in *lesv* mutants

Starch granule morphology (diameter and roundness) was analyzed using a granulomorphometer. A strong size reduction was observed in the *lesv* knockout mutant lines compared to WT plants whereas no change in granule size was observed in *esv1* mutants (Fig. 3). On average, a two-fold reduction in diameter was observed in *lesv* lines, with 50% of granules exhibiting a diameter between 4 µm and 5 µm in the mutants, compared to 13% in WT plants (Fig. 3a). This phenotype was consistently observed among all nine independent K.O. mutants. Additionally, the ellipsoid roundness of the granules was determined by measuring the ratio between the average radius of peripheral circles and the radius of the maximal circle included in the outlined particle (this ratio ranges from 0 to 1, with 1 corresponding to a perfect circle). Ellipsoid roundness was significantly decreased in the mutant lines, with an average roundness of 0.83 compared to 0.87 in WT plants (Fig. 3c). To further investigate the observed diameter reduction, purified starch granules or starch granules *in situ* were examined by optical or scanning electron microscopy (Fig. 4 and supplemental Fig. S5). In WT plants, the presence of large and regularly shaped granules was evidenced. In the *lesv* mutant lines, some granules of normal size could be observed. However, a strong accumulation of tiny starch granules was consistently seen in all the samples (Fig. 4 and supplemental Fig. S5). Atypical structures were also seen in the form of aggregates of fused small granules or forming composite structures that exhibited deep fissures (Fig. 4). Additionally, some of the small granules showed flat faces with sharp edges (Fig. 4). Differential Interference Contrast microscopy was employed to observe suspensions of purified starch granules submitted to ultrasonication and SDS washes to try separating aggregates (supplemental Fig. S5). Even when submitted to such treatments, aggregates did not disrupt suggesting glucan entanglement between granules.

**Figure 3:**
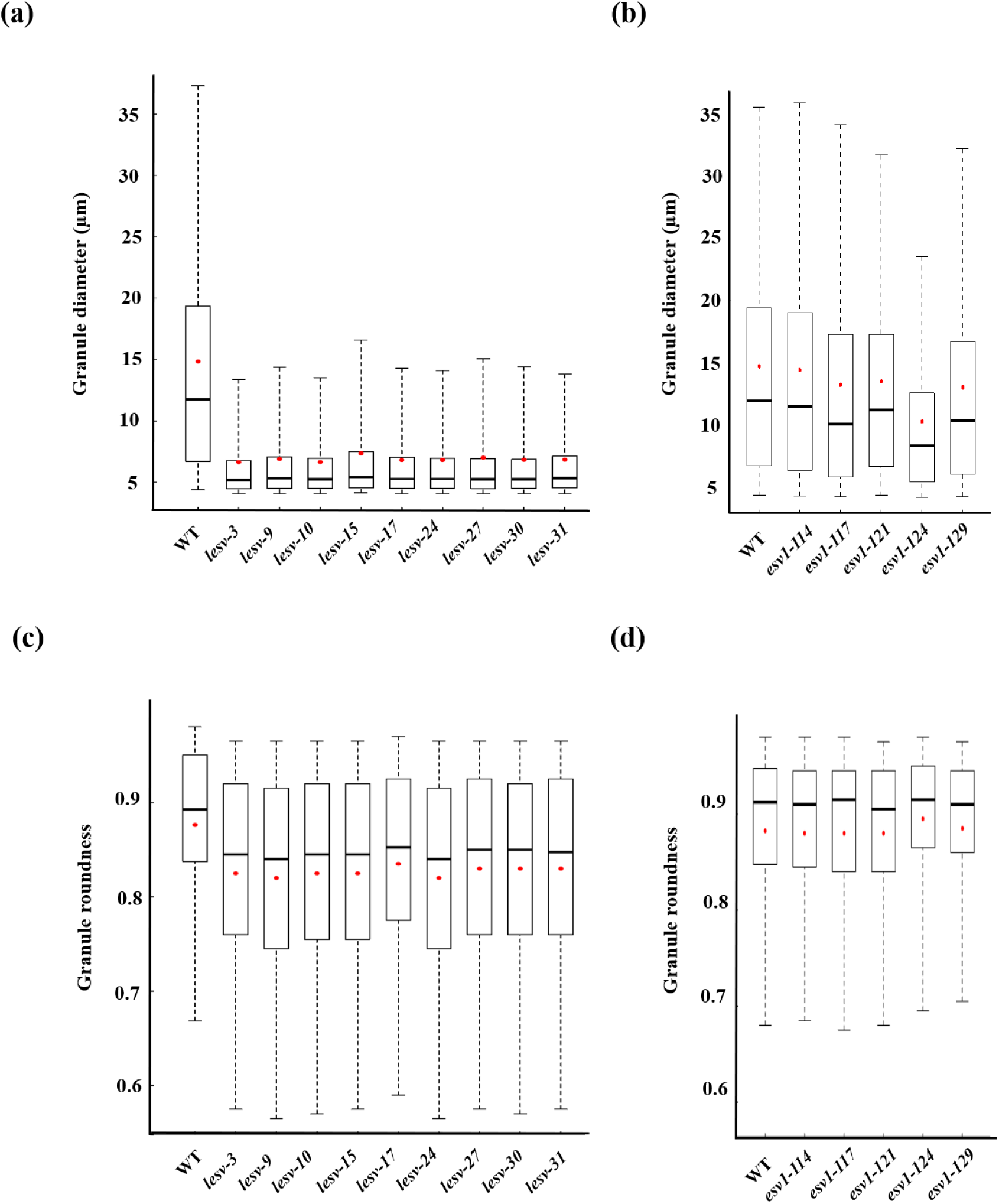
Starch granule granulomorphometry. (a, b) Starch granule diameter. (c, d) Starch granule roundness. For each measurement, the values corresponding to the percentiles P5, P25, P50, P75, and P95 were extracted from the dataset of each experimental replicate. The mean values were then calculated for each percentile across replicates of the same genotype. These averaged percentile values were used to generate the graphs. In the boxplots, the P5 and P95 values represent the extremities of the whiskers, the P25 and P75 values correspond to the bottom and top of the box, and the P50 (median) is shown as a black horizontal line within the box. Red dots represent the mean value for each mutant or wild-type line. For WT, data correspond to the mean of two biological replicates, each with two experimental replicates (n = 4). For mutant lines, data represent the mean of two experimental replicates (n = 2).

**Figure 4:**
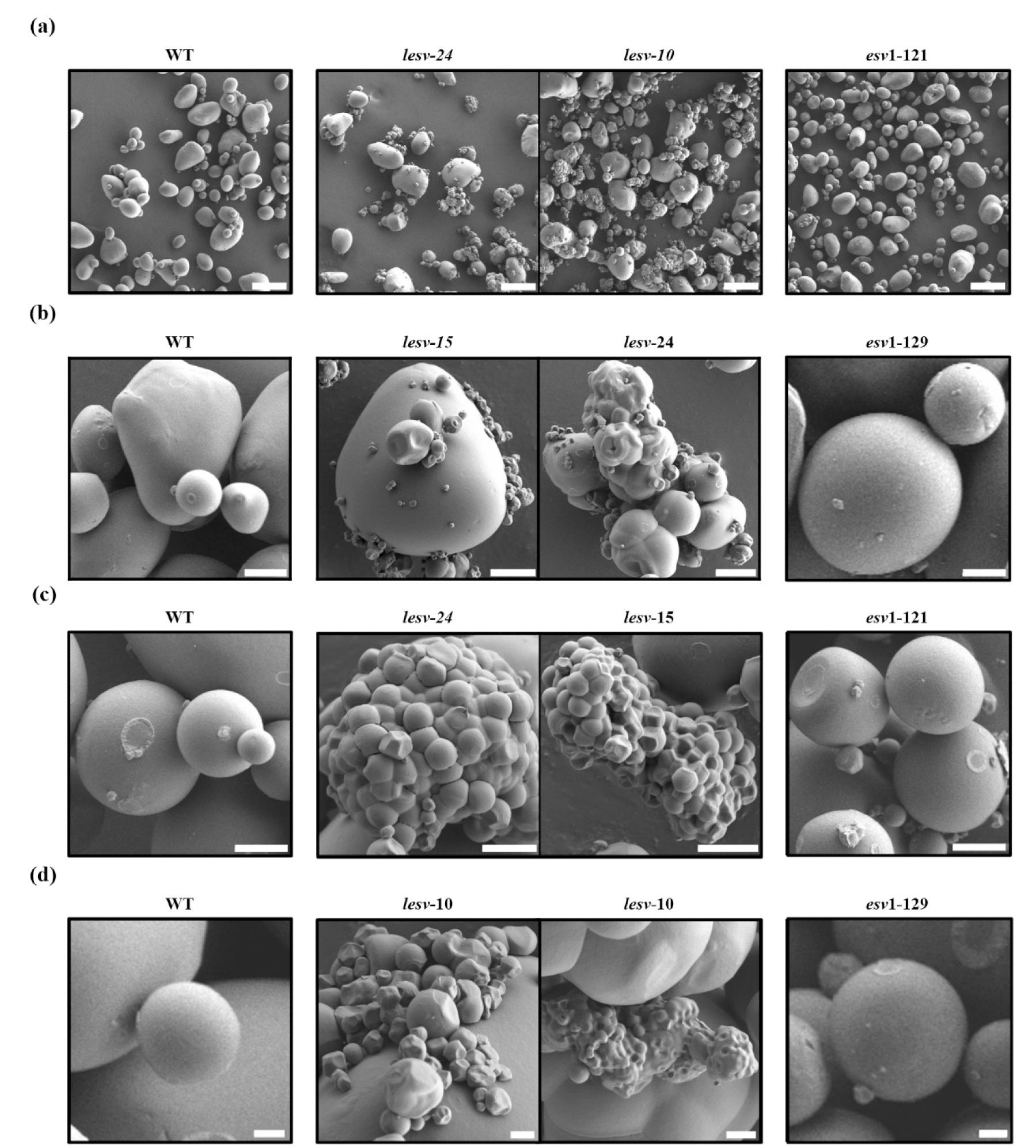
Scanning Electron Microscopy of purified starch granules of wild type (WT), *lesv* and *esv1* mutants. Starch samples were glued on an aluminum support with carbon double-sided tape and observed at 1 kV accelerating voltage with a Scanning Electron Microscope. Scale bar in (a) = 50µm; in (b) = 10µm; in (c) = 5µm and in (d) = 2µm.

### 3. Starch ultrastructure is unaffected in both *lesv* and *esv1* mutants

Starch composition and ultrastructure were investigated through several approaches (Fig. 5 and 6). Crystallinity was determined by Wide Angle X-ray Scattering (WAXS). All starch samples (WT and mutants) exhibited the same B-type structure showing a characteristic diffraction peak at 2θ = 7.4°. A slight decrease in crystallinity was observed in all the mutant lines, although not statistically significant (Fig. 5 and 6). Furthermore, the lamellar repeat distance (noted as “d”) was determined by Small Angle X-ray Scattering (SAXS) (Fig. 5 and 6). Results showed consistent d-value ranging between 7.2 and 7.4 for all starch samples. Radial orientation of starch glucan was confirmed by polarized light microscopy of starch suspensions (supplemental Fig. S3c). The birefringence cross was easily observed in both WT and the largest granules of *lesv* mutants. Its detection in the tiny granules aggregates was more challenging, although still discernible. In addition, amylose content (Fig. 5d and 6d) and the maximum wavelength of starch/iodine complex (Fig. 5c and 6c) were not significantly altered in the mutants compared to the WT. Moreover, the chain length distribution of amylopectin (CLD) was established, revealing no notable differences between the mutants and the WT (Fig. 5e, 5f, 6e and 6f).

**Figure 5:**
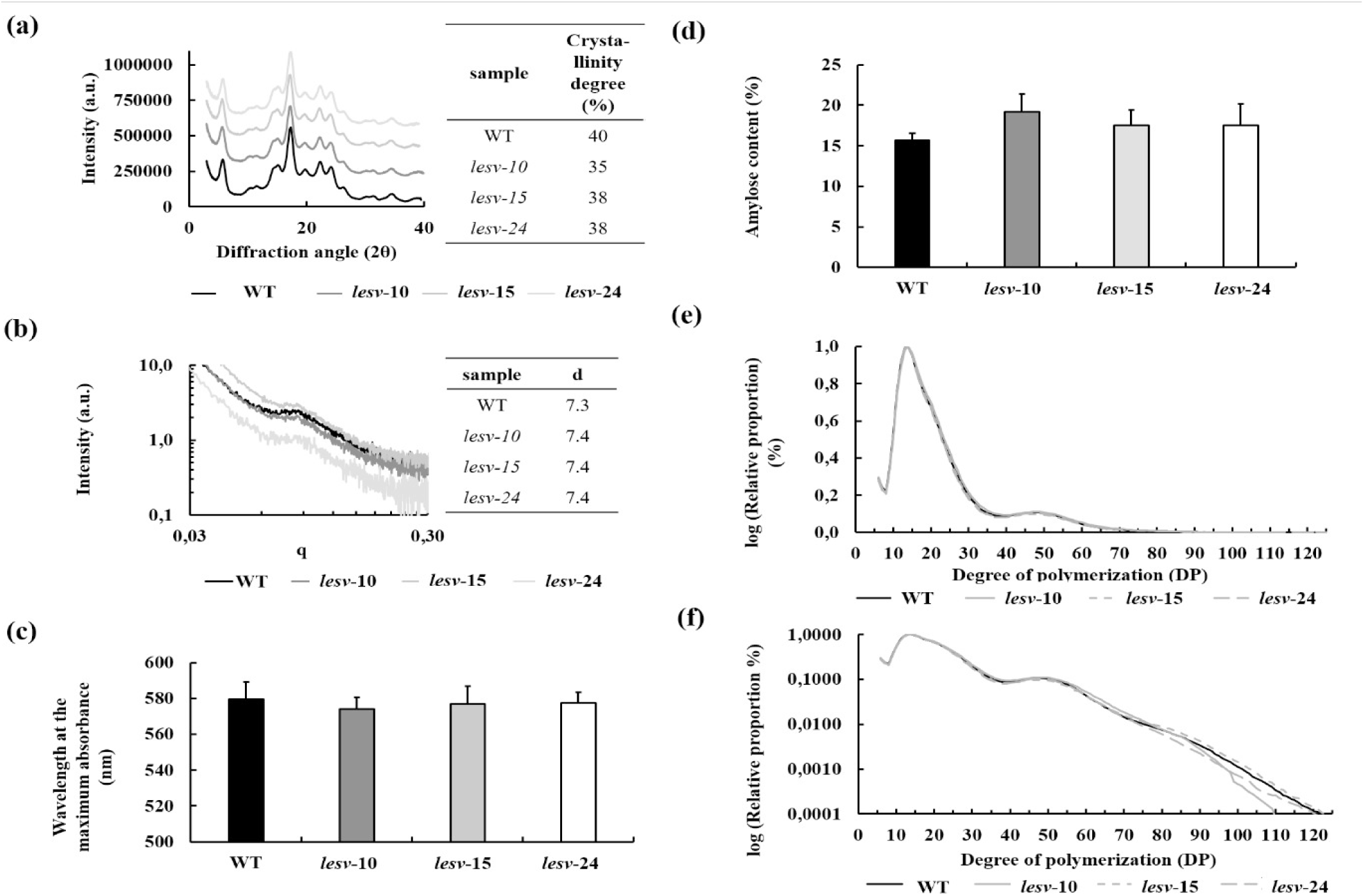
Molecular structure of *lesv* starches. (a) Crystalline-type determined by Wide Angle X-ray Scattering (WAXS). (b) Size of the lamellar repeat distance “d”, analyzed by Small Angle X-ray Scattering (SAXS). (c) Wavelength of the maximum absorbance of the iodine-starch complex of wild type (WT) and *lesv*. (d) Amylose content of WT and *lesv.* (e, f) Comparison of the chain length distribution (CLD) profiles of WT and *lesv* starch. Values were normalized according to the highest peak. For (a), (c) and (d), values are mean of three independent cultures. For (e) and (f) values are the mean (+/−SE) of three independent cultures and two experimental replicates. Statistical analysis was made using Mann-Whitney tests, comparing the mutant lines to the wild type.

**Figure 6:**
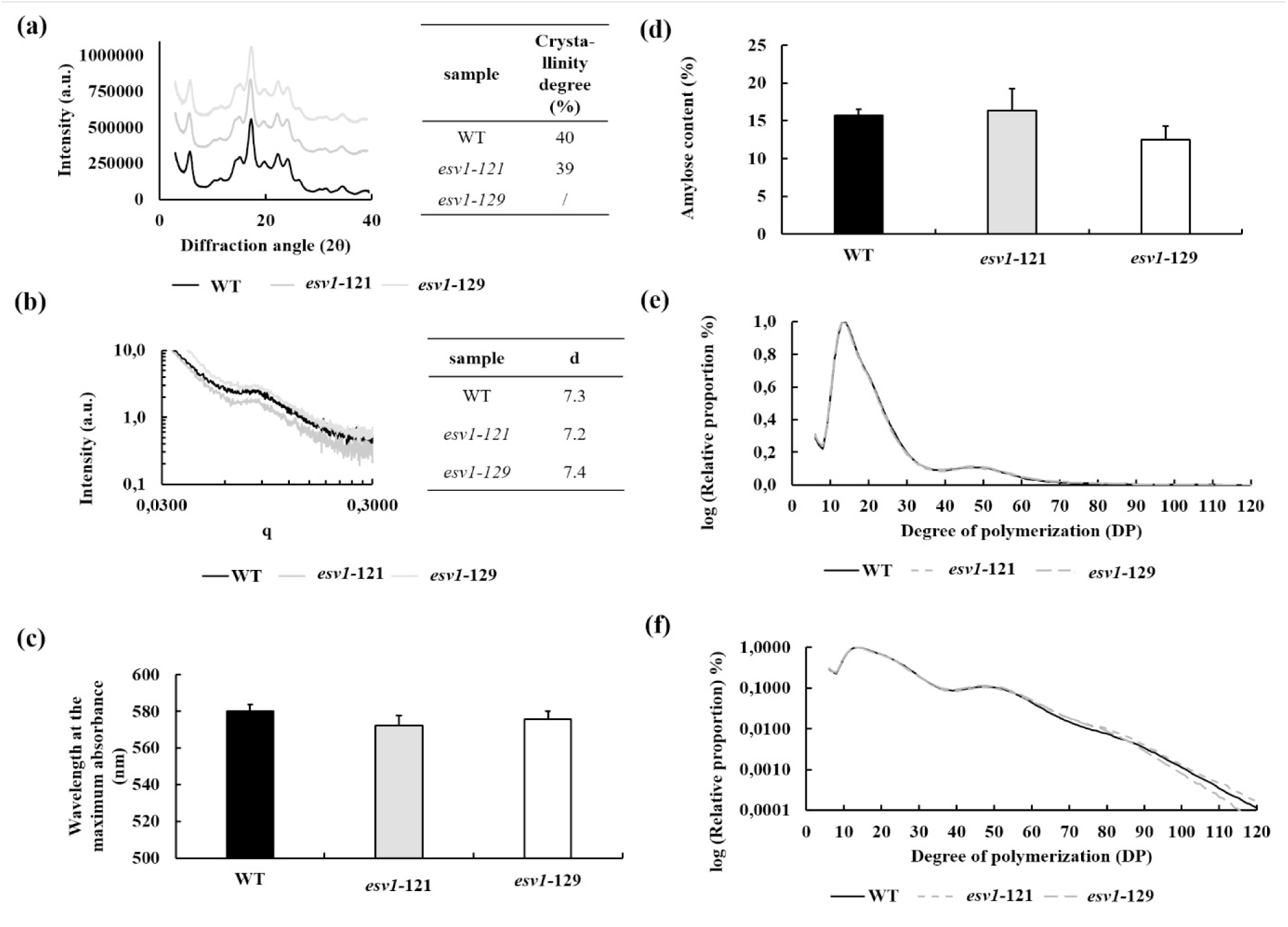
Molecular structure of *esv1* starches. (a) Crystalline-type determined by Wide Angle X-ray Scattering (WAXS). “/” indicates missing data. (b) Size of the lamellar repeat distance “d”, analyzed by Small Angle X-ray Scattering (SAXS). (c) Wavelength of the maximum absorbance of the iodine-starch complex of wild type (WT) and *esv1*. (d) Amylose content of WT and *esv1.* (e, f) Comparison of the chain length distribution (CLD) profiles of WT and *esv1* starch. Values were normalized according to the highest peak. For (a), (c) and (d), values are mean of three independent cultures. For (e) and (f) values are the mean (+/−SE) of three independent cultures and two experimental replicates. Statistical analysis was made using Mann-Whitney tests, comparing the mutant lines to the wild type.

### 4. *lesv* and *esv1* mutations affect starch phosphate content and α-amylolysis *in vitro*

To investigate the impact of *lesv* and *esv1* mutations on starch phosphorylation *in vivo*, phosphoester contents were determined by fluorescence-assisted capillary electrophoresis after acid hydrolysis (Fig. 7). A significant increase ranging from 19% to 28% of starch phosphorylation was measured in the *lesv* mutants. This increase was primarily attributed to an enrichment in O-6-phosphorylation, while the amount of O-3-phosphorylation was unchanged (Fig. 7a). This increase was also consistent with that already observed in fractionated potato starch granules of less than 10 µm diameter (Helle *et al*., 2019). In *esv1* mutant lines, a slight decrease of the phosphate content in the order of 18% to 20% was measured (Fig. 7b). The hydrolysis rate of mutant starches was assayed *in vitro* by measuring the increase of reducing ends after porcine pancreatic α-amylase treatment (Fig. 8). The degradability of *lesv* starch was significantly and consistently increased for all three tested mutant lines compared to WT (Fig. 8). On average, the *lesv* starch exhibited a hydrolysis rate of 26% after 48 hours, whereas that of the WT starch was only 17% representing an increase of the hydrolysis rate of 53%. This result was consistent with previous reports on the increased enzyme susceptibility of small granules due the higher amount of enzyme adsorption sites for a given starch weight (Noda *et al*., 2005; Dhital *et al*., 2010; Naguleswaran *et al*., 2012).

**Figure 7:**
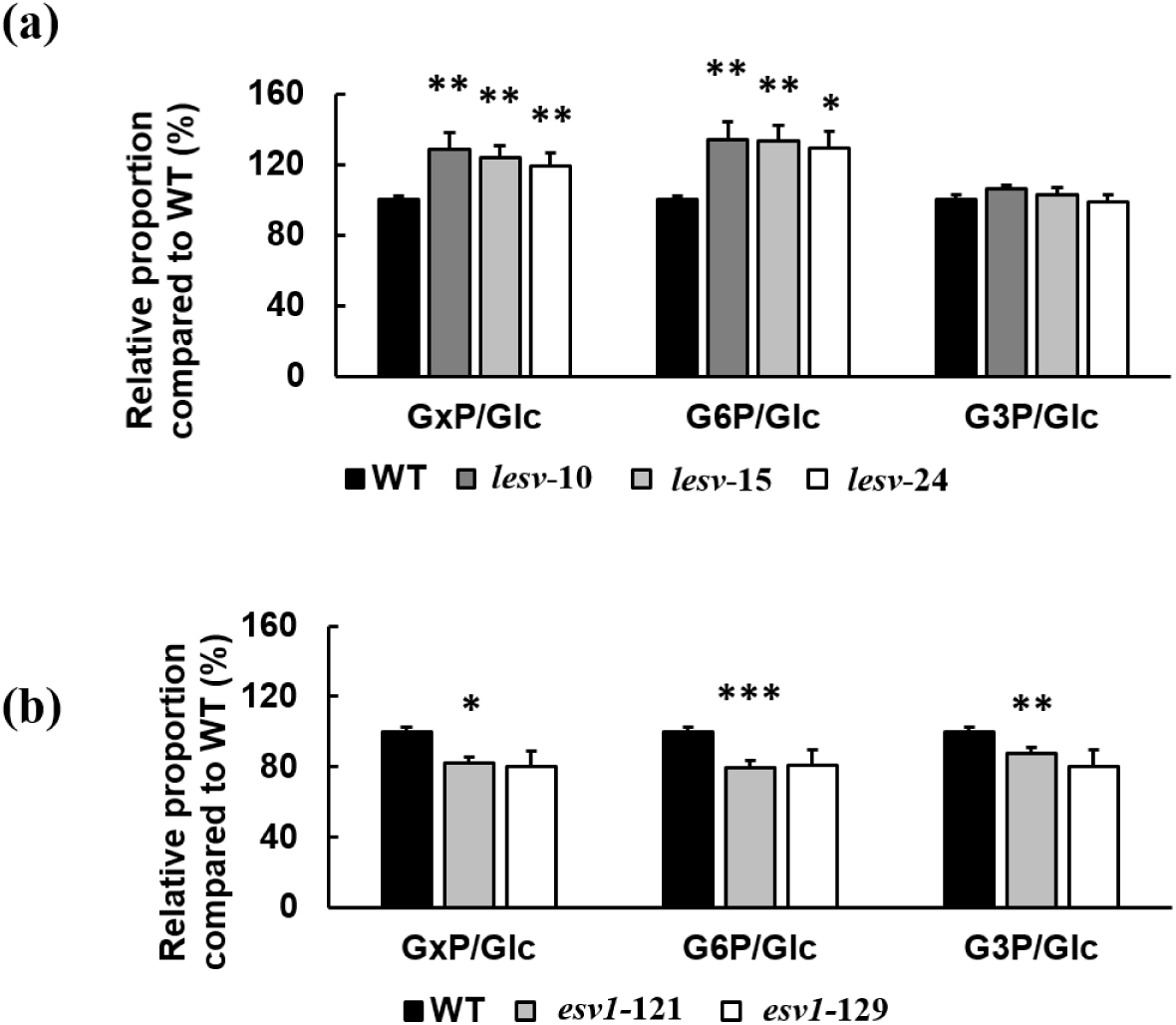
Assay of phosphoester residues: ratio of total phosphoester residues (GxP) over total glucose residues (GxP/Glc), ratio of O-6-phosphoesters (G6P/Glc) and ratio of O-3-phosphoesters (G3P/Glc) over total glucose residues in *lesv* (a) *and esv1* (b) mutants compared to wild-type (WT). Data were normalized according to the WT ratio (arbitrarily spotted at 100%). For WT, values are mean (+/− SE) of three biological replicates from three independent cultures and two experimental replicates for each culture (n=18). For each mutant lines, values are mean (+/− SE) of three independent cultures and two experimental replicates for each culture (n=6). Statistical analysis was made using Mann-Whitney tests, comparing the mutant line to the wild type (* = *p* ≤ 0,05; ** = *p* ≤ 0,01; *** = *p* ≤ 0,005).

**Figure 8:**
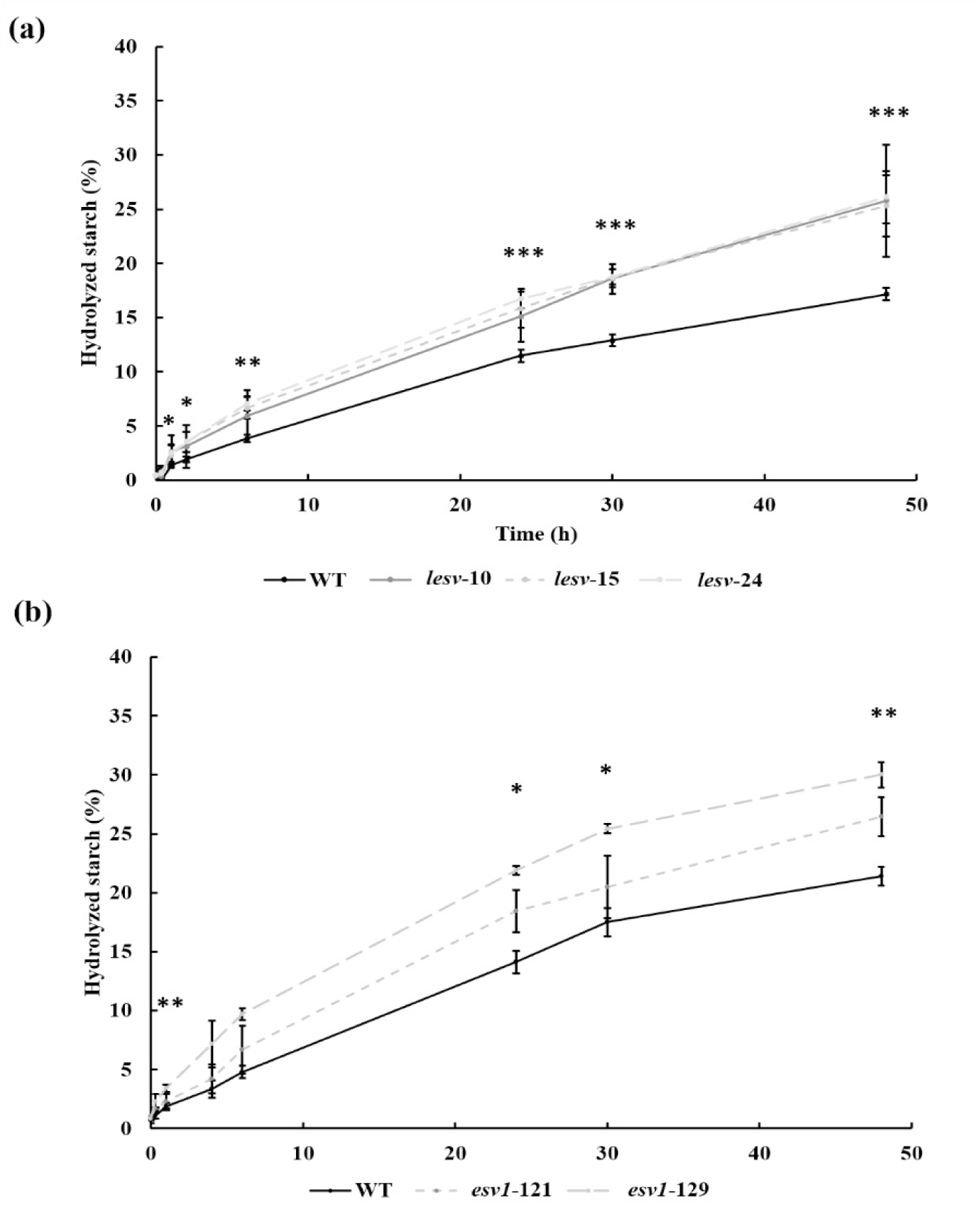
Enzymatic hydrolysis of starch by porcine pancreatic α-amylase (PPA). (a) *lesv* mutants compared to wild-type (WT) starch. (b) *esv1* mutants compared to WT starch. A suspension of 128U.mL^−1^ of PPA was added to ten milligrams of starch and incubated for 48h (x-axis) at 37°C. The % of hydrolyzed starch is plotted on the y-axis. For each line, starch was extracted from tubers of three independent cultures, each biological sample being submitted to three independent assays. Values are then mean (+/− SE) of these nine measures. Statistical analysis was made using Mann-Whitney tests, comparing the three mutant lines to the wild type (n=8-9; * = *p* ≤ 0,05; ** = *p* ≤ 0,01; *** = *p* ≤ 0,005).

### 5. LESV structural features and interaction with ISA1 support its role in regulating starch granule initiation

The alterations observed in *lesv* mutants suggest that LESV plays a key role in the initiation and growth of starch granules. Notably, the *lesv* phenotype closely resembled that of the *isa1* and *isa2* antisense lines, which also exhibited a marked reduction in granule size and the formation of aggregates (Bustos *et al*., 2004). Given that an interaction between LESV and ISA1 has been demonstrated in rice (Yan *et al*., 2024), we performed two independent yeast two-hybrid (Y2H) assays to confirm that this interaction is conserved in potato (Supplemental Fig. S6). These experiments consistently confirmed an interaction between potato LESV and ISA1. Additionally, the data suggest that ISA1 monomers can interact with each other, which is consistent with previous findings in maize and rice (Hussain *et al*., 2003; Utsumi & Nakamura, 2006; Kubo *et al*., 2010; Facon *et al*., 2013). To investigate structural differences between LESV and ESV1, we performed a protein sequence alignment using Clustal Omega, which revealed a distinctive N-terminal extension in LESV (Fig. 9a). Structural models generated with AlphaFold2 showed that both proteins share a conserved C-terminal domain composed of stacked antiparallel β-sheets (Fig. 9b), a feature also observed in Arabidopsis homologs (Osman *et al*., 2024). These β-sheets align tryptophan residues thought to mediate interactions with amylopectin. In contrast to ESV1, which possesses a short, disordered N-terminus, the N-terminal extension of LESV contains three predicted α-helices (Fig. 9c), supporting its unique functional role.

**Figure 9:**
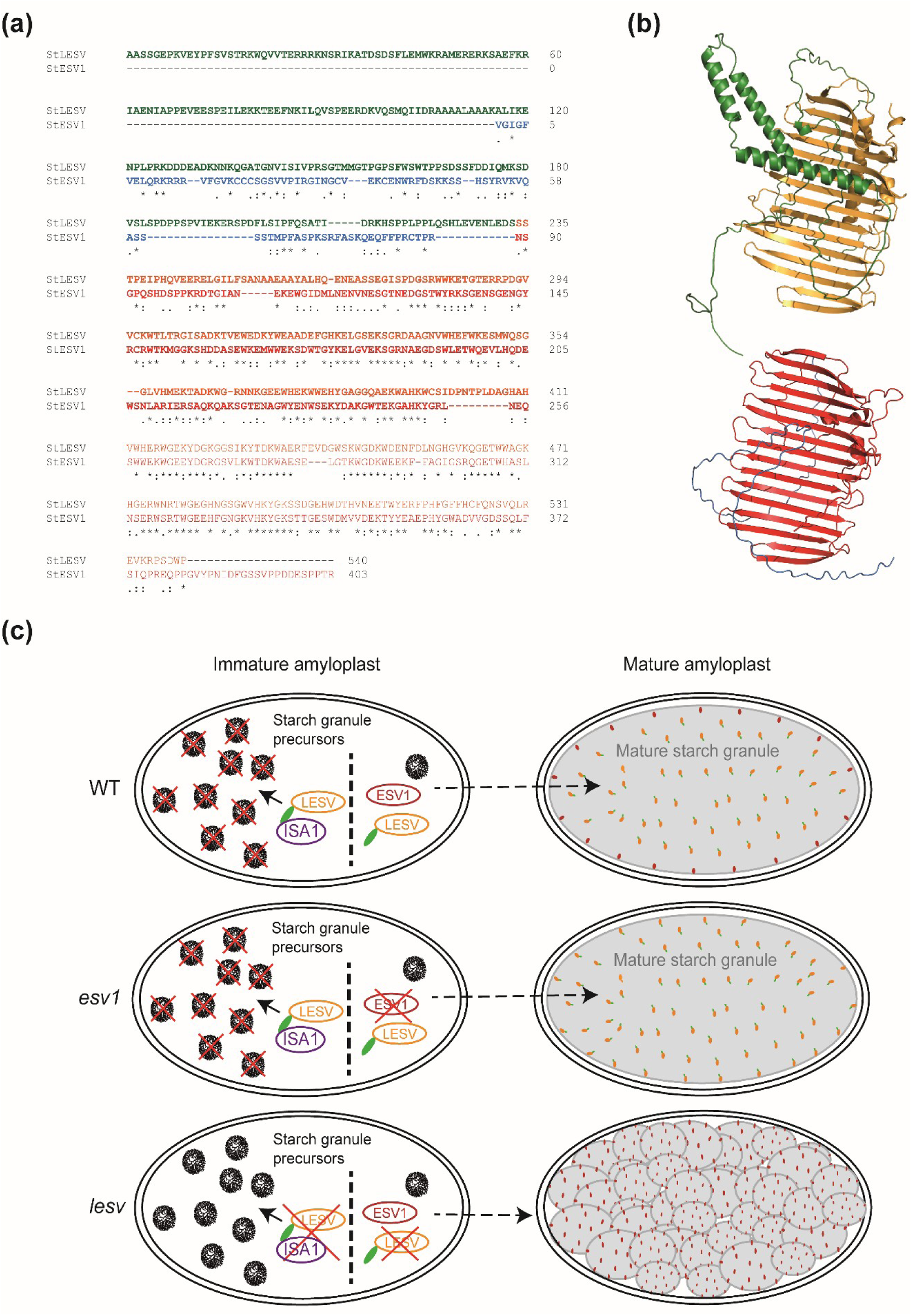
Proposed model for the functions of LESV and ESV1 in amyloplasts. **(a).** LESV and ESV1 protein sequences were retrieved from the potato genome sequence database and processed with Localizer software for transit peptide removal prior to alignment using Clustal omega. Orange and red lettering: conserved C-terminal domains. Green and blue lettering: highly divergent N-terminal regions of LESV and ESV1, respectively. **(b).** Protein models were generated with Colabfold software using Alphafold 2 and MMseqs2. The color code is the same as in (a). **(c).** Proposed model for LESV and ESV1 functions in potato amyloplasts. In wild-type plants (WT), LESV interacts with ISA1 (in purple) to regulate the number of soluble starch granule precursors. On the other hand, LESV and ESV1 interacts with glucans via their C-terminal domain to promote and stabilize phase transition, respectively, as proposed by (Liu *et al*., 2023), leading to the formation of a single starch granule in mature amyloplast. Inactivation of *ESV1* (middle panel) does not impact the number and structure of starch granules due to compensation by LESV. Inactivation of *LESV* deregulates the control of starch initiation and the promotion of phase transition is performed by ESV1. The abnormal high number of starch granule initiations leads to the synthesis of smaller starch granules and to granules aggregates.

## Discussion

### 1. *lesv* mutation alters the regulation of starch initiation in potato

The aim of this work was to investigate the function of the non-catalytic proteins LESV and ESV1 in potato tuber storage starch metabolism. To this end, K.O. mutants were generated by targeted genome editing using the CRISPR/Cas9 approach. Starch morphology was determined in nine or five independent K.O. lines for *LESV* or *ESV1*, respectively. For further in-depth phenotypic characterization, three or two K.O. lines were used, respectively. Interestingly, key agronomic traits, including plant growth, tuber and starch yield per plant, were unaffected by the mutations. However, the *lesv* mutation induced significant alterations in starch granule morphology, characterized by a significant reduction in diameter and in roundness, as determined by granulomorphometric analysis. Optical and electron microscopy confirmed the accumulation of numerous tiny granules, many of which were aggregated or fused structures. These aggregates were resistant to SDS and ultrasonication, suggesting tight glucan entanglement and the occurrence of multiple initiation events per amyloplast, characteristic of altered regulation of starch initiation. Strikingly, this phenotype closely mirrors that of *isa1* antisense lines previously described in potato (Bustos *et al*., 2004), where downregulation of isoamylase also led to small, aggregated starch granules. Phytoglycogen accumulation, a hallmark of isoamylase deficiency in *Arabidopsis* and cereals, was nonetheless measurable in *isa1* potato lines, albeit to a lesser extent, indicating species-specific differences in glucan fate following ISA1 deregulation. In addition to its function in starch deposition through amylopectin trimming, ISA1 was proposed to regulate starch initiation in potato tubers by limiting the number of initiation events.

Given the strong phenotypic overlap between *lesv* and *isa1* mutants, we hypothesized a functional interaction between LESV and ISA1. Our yeast two-hybrid (Y2H) assay confirmed that potato LESV physically interacts with ISA1, in line with recent findings in rice, where LESV (FLO9) was shown to bind ISA1 through its N-terminal domain (Yan et al., 2024). Furthermore, ISA1 self-interaction was also observed in our system, consistent with observations in maize and rice (Utsumi & Nakamura, 2006; Kubo et al., 2010; Facon et al., 2013). Despite high sequence similarity in their C-terminal regions, ESV1 and LESV seem to differ significantly in function. Both proteins possess a conserved C-terminal β-sheet domain with aligned tryptophan residues, known to bind starch (Feike et al., 2016; Osman et al., 2024). However, LESV has a unique N-terminal overhang, predicted to form α-helices and to undergo conformational changes upon binding amylopectin. This domain is absent in ESV1 and is required for ISA1 interaction in rice (Yan et al., 2024). Our results support this: potato *esv1* mutants showed no phenotype for granule morphology, suggesting that ESV1 lacks the capacity to recruit ISA1. A similar lack of phenotype was reported in *Arabidopsis esv1* mutants, although some irregularities in starch turnover were observed (Feike et al., 2016). Interestingly, the *lesv* phenotype in potato differs from the *lesv* phenotype in rice, as rice <u>lesv</u> mutants accumulate substantial amounts of phytoglycogen, contrary to the potato *lesv* mutants. This mirrors the difference in the phenotype of isa mutants: while rice *isa* mutants accumulate large amounts of phytoglycogen, potato mutants only show moderate phytoglycogen accumulation. Thus, although the nature of the phenotype differs between species, the fact that *lesv* mutants closely mimic *isa* mutants is conserved. This strengthens the case for a conserved functional interaction between LESV and ISA1 across species, despite divergent downstream effects.

We propose that in potato, LESV plays an essential role in regulating starch granule initiation through physical interaction with ISA1 and early glucan precursors (Fig. 9). In *lesv* mutants, this recruitment is lost, leading to deregulated initiation and formation of multiple granules per amyloplast. Meanwhile, ESV1 may still promote phase transition, maintaining crystalline structure, as starch ultrastructure remains unchanged. Despite the important differences between storage and transitory starch metabolism and their physiological functions in organs such as Arabidopsis leaves or potato tubers, the data reported here and the proposed model in Arabidopsis where LESV and ESV1 directly promote and stabilize glucan phase transition are not mutually exclusive. Although no major alteration in starch ultrastructure could be measured in potato *lesv* or *esv1* mutants, the loss of the corresponding proteins could induce slight changes that were not detected with the available methods. Finally, whether ESV1 has a redundant or supportive role in potato remains to be clarified; further analysis of double *lesv esv1* mutants will help address this.

### 2. *lesv* mutation affect starch phosphate content and α-amylolysis *in vitro*

The *lesv* mutation resulted in an increase in phosphate content, particularly due to elevated levels of O-6-phosphoester residues. This observation is consistent with two previous findings (Helle *et al*., 2019; Singh *et al*., 2022). First, it was shown that small starch granules in potato tuber exhibit higher phosphate content, mainly attributed to increased O-6-phosphoester residues (Helle *et al*., 2019). Moreover, *A. thaliana* LESV was found to regulate the phosphorylation activity of starch-related dikinases, key enzymes of the regulation of starch degradation (Singh *et al*., 2022). *In vitro* experiments revealed that LESV enhances the activity of PWD, an enzyme bound to starch granules involved in phosphorylating glucose at position C3 while reducing the activity of GWD which phosphorylated at position C6 crucial for PWD activity. The hydrolysis rate of *lesv* mutant starch is significantly increased compared to WT, showing a of 53% enhancement. The increased hydrolysis of *lesv* starch is consistent with previous findings. Smaller granules, due to their increased surface area per unit mass, facilitate enzyme diffusion and adsorption thereby, accelerating catalytic action (Dhital *et al*., 2010). This modification is particularly interesting as a perspective of industrial applications. The alteration of the digestibility is of economic interest, enabling the production of the same amount of hydrolysate in less time or with fewer enzymes. Additionally, small granules are valuable in composite materials, packaging components, fat replacers in low-fat food, wastewater treatment adsorbents, paper industry binders and as carriers for flavors, medicines or vitamins (Kim *et al*., 2015). While natural small granules from sources like quinoa, rice, amaranth and taro are typically used, producing small granules from potato could allow a higher yield production.

## Materials and Methods

### 1. Plant Material and Growth Conditions

The potato cultivar used in this study is *Solanum tuberosum* cv. Désirée. The genome of this cultivar has been resequenced by our laboratory, and a polymorphism map was generated at the genome scale (Sevestre *et al*., 2020). Plants were grown *in vitro* on MS medium (Murashig and Skoog) including vitamins (Duchefa), supplemented with 2% sucrose and 0.8% plant agar (Duchefa) in a growth cabinet under a 16 h light (80 µmol.m^−2^.s^−1^) : 8 h dark photoperiod, 22°C day : 20°C night. Plants were maintained and propagated *in vitro* by cuttings every six weeks.

### 2. Selection of Targets and Plasmids Constructs

Target sites adjacent to NGG protospacer adjacent motifs (PAM) were identified for *LESV* using Geneious software and CRISPOR online tool (http://crispor.tefor.net/) (Concordet & Haeussler, 2018). Single guide RNA (sgRNA) was selected according to predicted specificity, efficiency, out-of-frame scores, and Single Nucleotide Polymorphism (SNP) map (Sevestre *et al*., 2020). The vector used for mutagenesis is pDIRECT_22A (Addgene, plasmid #91133) (supplemental Fig. S3). It contains a kanamycin resistance gene. Plasmid vectors were constructed according to (Čermák *et al*., 2017). Briefly, oligonucleotides containing the sgRNA sequence were phosphorylated (for oligonucleotide sequences, see Table S1). By using a Golden Gate reaction, oligonucleotides were hybridized, creating a double strand DNA fragment with sticky ends complementary to AarI restriction site and introduced in the pDIRECT_22A vector digested by AarI. Plasmid construct was then multiplied by being produced in a chemically competent *Escherichia coli* TOP10 strain (genotype: FmcrA Δ(mrr-hsdRMS-mcrBC) Φ80lacZΔM15 ΔlacX74 recA1 araD139 Δ(ara-leu) 7697 galU galK rpsL (Str^R^) endA1 nupG λ-) (Invitrogen) and purified using the NucleoSpin Plasmid Kit of Macherey-Nagel.

### 3. *Agrobacterium*-mediated transformation

Agroinfection of potato was performed as described in (Sevestre *et al*., 2020). Briefly, *A. tumefaciens* strain GV3101, was transformed with 1 µg of the plasmid construct containing the sgRNA prior to selection on solid LB petri dishes containing 50 mg.L^−1^ of kanamycin as a selection agent. Potato explants of 3-4 mm in length from stems and petioles were prepared from three to four-week-old plants cultivated *in vitro*. They were placed on Petri dishes containing MS including vitamins (Duchefa) supplemented by 2% sucrose, 0.8% plant agar (Duchefa) and 1 mg.L^−1^ of indole-3-acetic acid (IAA), gibberellic acid (GA_3_) and zeatin-riboside (ZR) (Sigma Aldrich) for 48 h. Explants were then dipped into the *A. tumefaciens* culture with a DO_600_ adjusted to 0.2 using MS medium supplemented by 2% sucrose for 30 min with a slow mixing. Explants were then replaced on the same Petri dish for 48 h at 25°C in the dark. After removal of residual bacteria, explants were placed on Petri dishes containing MS medium including vitamins with 2% sucrose, 0.8% plant agar supplemented by 1 mg.L^−1^ of IAA, GA_3_ and ZR, 500 mg.L^−1^ of timentin and cefotaxime (Duchefa) and 50 mg.L^−1^ of kanamycin for two weeks to allow the development of calli. They were then placed on Petri dishes containing the same medium with an IAA concentration reduced to 0.1 mg.L^−1^ to allow the differentiation of the callus in stems. Those stems were taken and placed in tubes containing MS including vitamins with 2% sucrose and 0.8% plant agar and 50 mg.L^−1^ of kanamycin to allow the development of roots.

### 4. Characterization of Induced Mutations

DNA was extracted from leaves of three-week-old regenerated plants to assess the mutation caused by *Agrobacterium*-mediated transformation following the method by (Porebski *et al*., 1997). The presence of the Cas9 gene was checked by PCR using specific primers (Table S1) with GoTaq^®^ G2 Flexi DNA Polymerase (Promega). Primers specific to the targeted sequence were used to assess the Cas9-induced mutations (Table S1) and PCR products were analyzed by Sanger sequencing through subcontracting (Eurofins Genomics). Genotype was determined by analyzing the sequencing results using the ICE analysis tool by Synthego (https://ice.synthego.com/). K.O. mutants were selected according to R^2^ and K.O. scores.

### 5. Tuber Production and Starch Extraction

Three-week old *in vitro* plants were cultured in three liters pots in commercial soil. They were placed in a growth cabinet under a 16 h light (80 µmol.m^−2^.s^−1^):8 h dark photoperiod, 22°C day : 20°C night, for approximatively 12 weeks. After 4 weeks of culture, approximatively 5 cm of soil was added at the basis of the stem. Plants were watered twice a week for 11 weeks and tubers were harvested one week after the last watering, during plant senescence. Pictures of plants were taken every week. The background was removed on the plant and tubers pictures using the image editing tool removebg (https://www.remove.bg/). Three independent cultures were made, and the starch extracted from the tubers of these three replicates was used for the following experiments. After a washing step with tap water, tubers (100-120 g) were crushed on ice with deionized water using a Polytron homogenizer. Starch was filtered through a nylon net with a 100 µm mesh size (Sefar). Two to three centrifugation steps were performed at 4,000 g and 4°C using deionized water until water appears clean. The supernatant obtained from the first centrifugation step was collected, boiled at 100°C for 10 min and then stored at −20°C for subsequent measurements of the glucose and water-soluble polysaccharides content. Cellular debris were removed from starch through a centrifugation step at 4,000 g and 4°C for 45 min in a Percoll density gradient (Cytiva). The starch suspensions were then stored in 20% ethanol at 4°C for further analysis.

### 6. Starch, glucose and water-soluble polysaccharides content

To assay the starch content, starch suspensions were submitted to *in vitro* digestion with amyloglucosidase (AGS) from *Aspergillus niger* (Megazyme) for 30 min at 58°C. Subsequently, the starch content was measured using the Enzytec^TM^ Starch kit according to the manufacturer’s specifications. The content of the soluble extracts was also analyzed using the same kit, both before and after digestion with AGS, to measure the glucose and water-soluble polysaccharides contents, respectively.

### 7. Granulomorphometry

Starch granule size and shape were determined using a Flow-Cell FC200S+ granulomorphometer (Occhio). A filter was applied to exclude particles with a diameter smaller than 4 µm because of the camera resolution limits and a roundness inferior to 0.5 to exclude cellular debris. An average of 3,000 granules after filtration was analyzed for each sample. For determination of size and shape, the ISO area diameter and ellipsoid roundness parameters were retained.

### 8. Scanning electron microscopy

Purified starch granules were glued on an aluminum support with carbon double-sided tape and observed at 1 kV accelerating voltage with a Zeiss Merlin VP Scanning Electron Microscope.

### 9. Assay of Phosphorylated Glucose Residues in Starch

The quantification of phosphorylated glucose residues was made according to (Verbeke *et al*., 2016). Briefly, 1 mg of starch was hydrolyzed in 100 µL of 2 M trifluoroacetic acid (TFA) at 95°C for 80 min. 20 µL of the hydrolysate was added to 200 µL of ultrapure water and dried at room temperature in a speed-vac. After the addition of 200 µL of ultrapure water, a new drying step was performed. The samples were labeled using THF and APTS as described above. 96 µL of ultrapure water was added and 15 µL of this dilution was re-diluted in 185 µL of ultrapure water prior to injection in a Beckman Coulter PA800-plus Pharmaceutical Analysis System equipped with a laser induced fluorescence detector (Verbeke *et al*., 2016).

### 10. Wide-angle X-ray scattering

X-ray diffraction was used to identify the type of crystalline structure and to quantify the crystalline/amorphous structure ratio in the starch samples. Before analysis, starch water content was stabilized in a desiccator containing a saturated NaCl solution with a relative humidity of 75% for a week at room temperature. Fifty milligrams of sample was sealed in a copper ring between two adhesive tape sheets to prevent any change in water content during analysis. Two-dimensional diffraction diagrams were recorded using a Bruker D8 X-ray diffractometer (Karlsruhe, Germany) equipped with a GADDS detector. The X-ray radiation, Cu Kα1 (λ = 0.15406 nm), produced in a sealed tube at 40 kV and 40 mA, was selected and parallelized using crossed Göbel mirrors and collimated to produce a 500 µm beam diameter. Data were monitored by a GADDS 2D detector for 10 min and normalized.

### 11. Small-angle X-ray scattering (SAXS)

SAXS measurements were performed on a Xeuss 2.0 apparatus (Xenocs) equipped with point collimation (beam size: 300×300 μm2) and a micro source using a Cu-Kα radiation (λ =1.54 Å). The sample to detector distance was calibrated using silver behenate as a standard and was around 150 cm for the measurements. Through view 2D diffraction patterns were recorded on a Pilatus 200k detector (Dectris). From the 2D patterns integrated intensity profiles were computed using a Foxtrot® software. The lamellar repeat distance, designated as d, representing the alternative crystalline and amorphous lamellae, was calculated by the Bragg equation, *d = 2π/qmax*.

### 12. Maximum Wavelength of Starch/Iodine Complex

Two milligrams of starch were solubilized in 100% dimethylsulfoxide (DMSO) by heating 15 min at 100°C. After cooling down, they were diluted 100 times and 80 µL of the dilution was added to 20 µL of iodine solution (I_2_ 0.2% [w/v] and KI 2% [w/v]) and the absorbance was read between 500 and 700 nm using a SPECTROstar Nano spectrometer (BMG LABTECH’s), to determine the wavelength at the maximum of absorbance of the starch/iodine complex.

### 13. Determination of Amylose Content

To assess the amylose content in the samples, starch was fractionated using exclusion size chromatography to separate amylose and amylopectin molecules. 1.5 mg of starch was first gelatinized in 200 µL of dimethylsulfoxide (DMSO) by heating 15 min at 100°C, and precipitated in 800 µL of 100% ethanol to remove lipids. After 10 min of centrifugation at 10,000 g and 4°C, starch was then resolubilized in 300 µL of 10 mM sodium hydroxide. It was loaded on a column (0.5 cm x 65 cm), containing Sepharose CL-2B resin (Cytiva) equilibrated with 10mM sodium hydroxide. The flow rate was equilibrated at 12 mL.h^−1^. Fractions of 300 µL were collected. 80 µL of each fraction were mixed with 20 µL of iodine solution (I_2_ 0.2% [w/v] and KI 2% [w/v]) and analyzed by spectrophotometry between 500 and 700 nm. The fractions containing either amylose or amylopectin were pooled and the amount of each glucan was measured using the Enzytec^TM^ Starch kit.

### 14. Chain Length Distribution

The starch chain length distribution was measured according to (Helle *et al*., 2018). Briefly, two milligrams of starch were gelatinized in 250 µL of ultrapure water by heating at 100°C with regular stirring. It was then debranched at 42°C overnight with 20 U of isoamylase and 1 U of pullulanase (Sigma). The samples were then desalted on Alltech Carbograph columns and freeze-dried. Dried samples were labeled using 2 µL of cyanoborohydride in tetrahydrofuran (THF) and 2 µL of 200 mM 8-aminopyrene-1,3,6-trisulfonic acid (APTS) in 15% acetic acid and incubated at 42°C overnight. 46 µL of ultrapure water was added and the samples were diluted 50 times with ultrapure water. They were then analyzed in a Beckman Coulter PA800-plus Pharmaceutical Analysis System equipped with a laser induced fluorescence detector according to (Fermont *et al*., 2022).

### 15. *In vitro* enzymatic hydrolysis of starch

The enzymatic degradation of *lesv* starch was assayed by measuring the production of reducing glucose using 3,5-dinitrosalicylic acid (DNS). The enzyme used to hydrolyze starch is the porcine pancreatic α-amylase (PPA) (Megazyme). 128 U/mL of PPA was suspended in a 20 mM phosphate, 0.02 % sodium azide, 2 mM NaCl, 0.25 mM CaCl_2_ pH 7.0 buffer. 1 mL of the enzyme suspension was added to 10 mg of starch in a 1.5 mL Eppendorf. The mixture was incubated at 37°C in a thermomixer and stirred. The estimation of hydrolyzed starch was performed by taking 15 µL aliquots after 0, 20, 60, 360, 1440, 1800 and 2880 min and mixing it with a DNS solution, according to (Miller, 1959). After heating 10 min at 37°C, the absorbance was read at 540 nm in a spectrophotometer. Standard curves of glucose were realized (0, 1, 2, 3, 4 and 5 g.L^−1^). A negative control containing no PPA was realized for each sample and subtracted to the measures.

### 16. Yeast two-hybrid assay

The coding sequences of *LESV* and *ISA1* genes were ordered as a gBlocks DNA fragment flanked by *attB* Gateway recombination sites. These sequences were cloned into the Gateway entry vector pDONR221 using the BP Clonase II enzyme kit (Thermo Scientific). The resulting entry clones were subsequently recombined into the destination vectors pGADT7 (containing the activation domain) or pGBKT7 (containing the DNA-binding domain) via LR Clonase II-mediated recombination (Thermo Scientific). Various plasmid combinations were co-transformed into *Saccharomyces cerevisiae* strain AH109 (*MATa, trp1-901, leu2-3, 112, ura3-52, his3-200, gal4Δ, gal80Δ, LYS2::GAL1UAS-GAL1TATA-HIS3, MEL1 GAL2UAS-GAL2TATA-ADE2, URA3::MEL1UAS-MEL1TATA-lacZ*). Transformed yeast cells were selected on Double Dropout medium (DDO, SD/- Leu-Trp) (Takara) to confirm the presence of both plasmids and screened for protein-protein interactions on Quadruple Dropout medium (QDO, SD/-Leu-Trp-Ade-His) (Takara), where growth indicates a positive interaction.

### 17. Protein alignment and modelling

Protein sequences were retrieved from the potato genome assembly dataset (v6.1) for the doubled monoploid potato DM 1-3 516 R44 (Pham *et al*., 2020). Plastid transit peptides were predicted with the use of Localizer (https://localizer.csiro.au). Protein sequences were aligned using clustal omega (Madeira *et al*., 2024). ESV1 or LESV sequences without transit peptide were then computed with Colabfold v1.5.5 software based on AlphaFold2 and MMseqs2 (Jumper *et al*., 2021; Mirdita *et al*., 2022). Colabfold was executed with default parameters, and five structural models were generated per input sequence. Models were ranked based on AlphaFold’s predicted local distance difference test (pLDDT) score. The highest-confidence model (pLDDT ≥ 70) was selected for figures.

## Acknowledgments

This research was supported by ANR grants through the JCJC and PRC programs (ANR-17-CE20-0012 and ANR-22-CE44-0010, respectively) and the Hauts-de-France region through the CPER “Alibiotech”, as well as the ANRT through the Ph.D of CL (ANRT n° 2022/0449). DS is supported by BBSRC grant BB/X01097X/1. The authors thank Corentin Spriet and Oceane George for assistance with optical microscopy and tuber starch extraction, respectively, Nicolas Barois of US 41 - UAR 2014 - PLBS for SEM imaging and Lèna Brionne for assistance with starch α-amylolysis experiments. Financial support from Région Nord Pas-de-Calais and European FEDER for the SAXS laboratory equipment is gratefully acknowledged.

## References

Ball S, Guan H-P, James M, Myers A, Keeling P, Mouille G, Buléon A, Colonna P, Preiss J. 1996. From Glycogen to Amylopectin: A Model for the Biogenesis of the Plant Starch Granule. Cell 86: 349–352.

Ball SG, Van De Wal MHBJ, Visser RGF. 1998. Progress in understanding the biosynthesis of amylose. Trends in Plant Science 3: 462–467.

Bustos R, Fahy B, Hylton CM, Seale R, Nebane NM, Edwards A, Martin C, Smith AM. 2004. Starch granule initiation is controlled by a heteromultimeric isoamylase in potato tubers. Proceedings of the National Academy of Sciences 101: 2215–2220.

Čermák T, Curtin SJ, Gil-Humanes J, Čegan R, Kono TJY, Konečná E, Belanto JJ, Starker CG, Mathre JW, Greenstein RL, et al. 2017. A Multipurpose Toolkit to Enable Advanced Genome Engineering in Plants. The Plant Cell 29: 1196–1217.

Concordet J-P, Haeussler M. 2018. CRISPOR: intuitive guide selection for CRISPR/Cas9 genome editing experiments and screens. Nucleic Acids Research 46: W242–W245.

Delatte T, Trevisan M, Parker ML, Zeeman SC. 2005. Arabidopsis mutants *Atisa1* and *Atisa2* have identical phenotypes and lack the same multimeric isoamylase, which influences the branch point distribution of amylopectin during starch synthesis. The Plant Journal 41: 815–830.

Dhital S, Shrestha AK, Gidley MJ. 2010. Relationship between granule size and in vitro digestibility of maize and potato starches. Carbohydrate Polymers 82: 480–488.

Facon M, Lin Q, Azzaz AM, Hennen-Bierwagen TA, Myers AM, Putaux J-L, Roussel X, D’Hulst C, Wattebled F. 2013. Distinct functional properties of isoamylase-type starch debranching enzymes in monocot and dicot leaves. Plant Physiology 163: 1363–1375.

Feike D, Seung D, Graf A, Bischof S, Ellick T, Coiro M, Soyk S, Eicke S, Mettler-Altmann T, Lu KJ, et al. 2016. The Starch Granule-Associated Protein EARLY STARVATION1 Is Required for the Control of Starch Degradation in *Arabidopsis thaliana* Leaves. The Plant Cell 28: 1472–1489.

Fermont L, Szydlowski N, Colleoni C. 2022. Determination of Glucan Chain Length Distribution of Glycogen Using the Fluorophore-Assisted Carbohydrate Electrophoresis (FACE) Method. Journal of Visualized Experiments: 63392.

Fredriksson H, Silverio J, Andersson R, Eliasson A-C, Åman P. 1998. The influence of amylose and amylopectin characteristics on gelatinization and retrogradation properties of different starches. Carbohydrate Polymers 35: 119–134.

Helle S, Bray F, Putaux J-L, Verbeke J, Flament S, Rolando C, D’Hulst C, Szydlowski N. 2019. Intra-Sample Heterogeneity of Potato Starch Reveals Fluctuation of Starch-Binding Proteins According to Granule Morphology. Plants 8: 324.

Helle S, Bray F, Verbeke J, Devassine S, Courseaux A, Facon M, Tokarski C, Rolando C, Szydlowski N. 2018. Proteome Analysis of Potato Starch Reveals the Presence of New Starch Metabolic Proteins as Well as Multiple Protease Inhibitors. Frontiers in Plant Science 9: 746.

Hussain H, Mant A, Seale R, Zeeman S, Hinchliffe E, Edwards A, Hylton C, Bornemann S, Smith AM, Martin C, et al. 2003. Three Isoforms of Isoamylase Contribute Different Catalytic Properties for the Debranching of Potato Glucans[W]. The Plant Cell 15: 133–149.

Jumper J, Evans R, Pritzel A, Green T, Figurnov M, Ronneberger O, Tunyasuvunakool K, Bates R, Žídek A, Potapenko A, et al. 2021. Highly accurate protein structure prediction with AlphaFold. Nature 596: 583–589.

Jung Y-J, Nogoy FM, Lee S-K, Cho Y-G, Kang K-K. 2018. Application of ZFN for Site Directed Mutagenesis of Rice SSIVa Gene. Biotechnology and Bioprocess Engineering 23: 108–115.

Kim H-Y, Park SS, Lim S-T. 2015. Preparation, characterization and utilization of starch nanoparticles. Colloids and Surfaces B: Biointerfaces 126: 607–620.

Kram AM, Oostergetel GT, Van Burggen E. 1993. Localization of Branching Enzyme in Potato Tuber Cells with the Use of lmmunoelectron Microscopy. 101: 237–243.

Kubo A, Colleoni C, Dinges JR, Lin Q, Lappe RR, Rivenbark JG, Meyer AJ, Ball SG, James MG, Hennen-Bierwagen TA, et al. 2010. Functions of heteromeric and homomeric isoamylase-type starch-debranching enzymes in developing maize endosperm. Plant Physiology 153: 956–969.

Liu C, Pfister B, Osman R, Ritter M, Heutinck A, Sharma M, Eicke S, Fischer-Stettler M, Seung D, Bompard C, et al. 2023. LIKE EARLY STARVATION 1 and EARLY STARVATION 1 promote and stabilize amylopectin phase transition in starch biosynthesis. Science Advances 9: eadg7448.

Lu K-J, Pfister B, Jenny C, Eicke S, Zeeman SC. 2018. Distinct Functions of STARCH SYNTHASE 4 Domains in Starch Granule Formation. Plant Physiology 176: 566–581.

Madeira F, Madhusoodanan N, Lee J, Eusebi A, Niewielska A, Tivey ARN, Lopez R, Butcher S. 2024. The EMBL-EBI Job Dispatcher sequence analysis tools framework in 2024. Nucleic Acids Research 52: W521–W525.

Mérida A, Fettke J. 2021. Starch granule initiation in *Arabidopsis thaliana* chloroplasts. The Plant Journal 107: 688–697.

Miller GL. 1959. Use of Dinitrosalicylic Acid Reagent for Determination of Reducing Sugar. Analytical Chemistry 31: 426–428.

Mirdita M, Schütze K, Moriwaki Y, Heo L, Ovchinnikov S, Steinegger M. 2022. ColabFold: making protein folding accessible to all. Nature Methods 19: 679–682.

Naguleswaran S, Li J, Vasanthan T, Bressler D, Hoover R. 2012. Amylolysis of large and small granules of native triticale, wheat and corn starches using a mixture of α-amylase and glucoamylase. Carbohydrate Polymers 88: 864–874.

Noda T, Takigawa S, Matsuura-Endo C, Kim S-J, Hashimoto N, Yamauchi H, Hanashiro I, Takeda Y. 2005. Physicochemical properties and amylopectin structures of large, small, and extremely small potato starch granules. Carbohydrate Polymers 60: 245–251.

Osman R, Bossu M, Dauvillée D, Spriet C, Liu C, Zeeman SC, D’Hulst C, Bompard C. 2024. LIKE EARLY STARVATION 1 interacts with amylopectin during starch biosynthesis. Plant Physiology: kiae193.

Pham GM, Hamilton JP, Wood JC, Burke JT, Zhao H, Vaillancourt B, Ou S, Jiang J, Buell CR. 2020. Construction of a chromosome-scale long-read reference genome assembly for potato. GigaScience 9: giaa100.

Porebski S, Bailey LG, Baum BR. 1997. Modification of a CTAB DNA extraction protocol for plants containing high polysaccharide and polyphenol components. Plant Molecular Biology Reporter 15: 8–15.

Reyniers S, Ooms N, Gomand SV, Delcour JA. 2020. What makes starch from potato (*Solanum tuberosum* L.) tubers unique: A review. Comprehensive Reviews in Food Science and Food Safety 19: 2588–2612.

Roldán I, Wattebled F, Mercedes Lucas M, Delvallé D, Planchot V, Jiménez S, Pérez R, Ball S, D’Hulst C, Mérida Á. 2007. The phenotype of soluble starch synthase IV defective mutants of *Arabidopsis thaliana* suggests a novel function of elongation enzymes in the control of starch granule formation. The Plant Journal 49: 492–504.

Satoh H, Shibahara K, Tokunaga T, Nishi A, Tasaki M, Hwang S-K, Okita TW, Kaneko N, Fujita N, Yoshida M, et al. 2008. Mutation of the Plastidial α-Glucan Phosphorylase Gene in Rice Affects the Synthesis and Structure of Starch in the Endosperm. The Plant Cell 20: 1833–1849.

Seung D, Boudet J, Monroe J, Schreier TB, David LC, Abt M, Lu K-J, Zanella M, Zeeman SC. 2017. Homologs of PROTEIN TARGETING TO STARCH Control Starch Granule Initiation in Arabidopsis Leaves. The Plant Cell 29: 1657–1677.

Seung D, Lu K-J, Stettler M, Streb S, Zeeman SC. 2016. Degradation of Glucan Primers in the Absence of Starch Synthase 4 Disrupts Starch Granule Initiation in Arabidopsis. Journal of Biological Chemistry 291: 20718–20728.

Seung D, Schreier TB, Bürgy L, Eicke S, Zeeman SC. 2018. Two Plastidial Coiled-Coil Proteins Are Essential for Normal Starch Granule Initiation in Arabidopsis. The Plant Cell 30: 1523–1542.

Seung D, Smith AM. 2019. Starch granule initiation and morphogenesis—progress in Arabidopsis and cereals. Journal of Experimental Botany 70: 771–784.

Sevestre F, Facon M, Wattebled F, Szydlowski N. 2020. Facilitating gene editing in potato: a Single-Nucleotide Polymorphism (SNP) map of the Solanum tuberosum L. cv. Desiree genome. Scientific Reports 10: 2045.

Sharma M, Abt MR, Eicke S, Ilse TE, Liu C, Lucas MS, Pfister B, Zeeman SC. 2024. MFP1 defines the subchloroplast location of starch granule initiation. Proceedings of the National Academy of Sciences 121: e2309666121.

Singh A, Compart J, AL-Rawi SA, Mahto H, Ahmad AM, Fettke J. 2022. LIKE EARLY STARVATION 1 alters the glucan structures at the starch granule surface and thereby influences the action of both starch-synthesizing and starch-degrading enzymes. The Plant Journal 111: 819–835.

Streb S, Delatte T, Umhang M, Eicke S, Schorderet M, Reinhardt D, Zeeman SC. 2009. Starch Granule Biosynthesis in *Arabidopsis* Is Abolished by Removal of All Debranching Enzymes but Restored by the Subsequent Removal of an Endoamylase. The Plant Cell 20: 3448–3466.

Toyosawa Y, Kawagoe Y, Matsushima R, Crofts N, Ogawa M, Fukuda M, Kumamaru T, Okazaki Y, Kusano M, Saito K, et al. 2016. Deficiency of Starch Synthase IIIa and IVb Alters Starch Granule Morphology from Polyhedral to Spherical in Rice Endosperm. Plant Physiology 170: 1255–1270.

Utsumi Y, Nakamura Y. 2006. Structural and enzymatic characterization of the isoamylase1 homo-oligomer and the isoamylase1–isoamylase2 hetero-oligomer from rice endosperm. Planta 225: 75–87.

Vandromme C, Spriet C, Dauvillée D, Courseaux A, Putaux J, Wychowski A, Krzewinski F, Facon M, D’Hulst C, Wattebled F. 2019. PII1: a protein involved in starch initiation that determines granule number and size in Arabidopsis chloroplast. New Phytologist 221: 356–370.

Vandromme C, Spriet C, Putaux J, Dauvillée D, Courseaux A, D’Hulst C, Wattebled F. 2023. Further insight into the involvement of PII1 in starch granule initiation in Arabidopsis leaf chloroplasts. New Phytologist 239: 132–145.

Verbeke J, Penverne C, D’Hulst C, Rolando C, Szydlowski N. 2016. Rapid and sensitive quantification of C3- and C6-phosphoesters in starch by fluorescence-assisted capillary electrophoresis. Carbohydrate Polymers 152: 784–791.

Vos-Scheperkeuter GH, De Boer W, Visser RGF, Feenstra WJ, Witholt B. 1986. Identification of Granule-Bound Starch Synthase in Potato Tubers. Plant Physiology 82: 411–416.

Wattebled F, Dong Y, Dumez S, Delvallé D, Planchot V, Berbezy P, Vyas D, Colonna P, Chatterjee M, Ball S, et al. 2005. Mutants of Arabidopsis Lacking a Chloroplastic Isoamylase Accumulate Phytoglycogen and an Abnormal Form of Amylopectin. Plant Physiology 138: 184–195.

Wattebled F, Planchot V, Dong Y, Szydlowski N, Pontoire B, Devin A, Ball S, D’Hulst C. 2008. Further Evidence for the Mandatory Nature of Polysaccharide Debranching for the Aggregation of Semicrystalline Starch and for Overlapping Functions of Debranching Enzymes in Arabidopsis Leaves. Plant Physiology 148: 1309–1323.

Yan H, Zhang W, Wang Y, Jin J, Xu H, Fu Y, Shan Z, Wang X, Teng X, Li X, et al. 2024. Rice LIKE EARLY STARVATION1 cooperates with FLOURY ENDOSPERM6 to modulate starch biosynthesis and endosperm development. The Plant Cell: koae006.

